# Reconstruction of Bone Defect Based on the Fusion of Osteoblasts and MSM/HA/PLGA Porous Scaffolds

**DOI:** 10.1101/2020.04.15.042713

**Authors:** Xiaodong Wu, Yanchen Zheng, Jian Chen, Qiang Xu, Pengpeng Li, Conghui Han, Weiling Huo

## Abstract

Reconstruction of bone defect is one of the difficult problems in orthopedic treatment, and bone tissue scaffold implantation is the most promising direction of bone defect reconstruction.

In this study, we used the combination of HA (Hydroxyapatite) and PLGA [Poly (lactic-co-glycolic acid)] in the construction of polymer scaffolds, and introduced bioactive MSM (Methyl sulfonyl methane) into polymer scaffolds to prepare porous scaffolds. The osteoblasts, isolated and cultured in vitro, were seeded in the porous scaffolds to construct tissue-engineered scaffolds. Meanwhile, the model of rabbit radius defect was constructed to evaluate the biological aspects of five tissue-engineered scaffolds, which provided experimental basis for the application of the porous scaffolds in bone tissue engineering.

The SEM characterization showed the pore size of porous scaffolds was uniform and the porosity was about 90%. The results of contact Angle testing suggested that the hydrophobic porous scaffold surface could effectively promote cell adhesion and cell proliferation, while mechanical property test showed good machinability. The results of drug loading and release efficiency of MSM showed that porous scaffolds could load MSM efficiently and prolong the release time of MSM.

In vitro incubation of porous scaffolds and osteoblasts showed that the addition of a small quantity of MSM could promote the infiltration and proliferation of osteoblasts on the porous scaffolds. Similar results were obtained by implanting the tissue-engineered scaffolds, fused with the osteoblasts and MSM/HA/PLGA porous scaffolds, into the rabbit radius defect, which provided experimental basis for the application of the MSM/HA/PLGA porous scaffolds in bone tissue engineering.

## Introduction

Bone defects are bone shortages caused by trauma or surgery that often result in bone disjunction, delayed healing, and local dysfunction. Bone defect is a common disease in clinic, and bone defect reconstruction is one of the difficult problems in orthopedic treatment[1, 2].

At present, many methods are used to repair bone defects, including transplantation of autologous or allogeneic bone [1], stromal stem cells [3], and tissue engineering scaffold repairing. Bone transplantation has been widely used in clinical practice, with good results. However, this repair strategy still has the risk of secondary surgery, transplant rejection and infection of disease. It is also limited by the lack of autologous bone resources, allograft rejection and other factors. In order to overcome these risks and limitations of bone transplantation, Tissue engineering scaffolds have been studied all over the world.

Tissue engineering scaffold refers to the material that can be implanted into an organism and can be combined with living tissue cells to realize tissue functions. The Ideal scaffold material can simulate the characteristics of the natural extracellular matrix (ECM) to promote cell adhesion, proliferation and differentiation.

The ideal bone tissue scaffold should have the following characteristics: good bone conductivity and bone inductivity; appropriate aperture and porosity; certain mechanical strength and plasticity. The materials used to prepare bone scaffolds include degradable inorganic materials, degradable polymer materials, non-biodegradable materials and composite materials, while composite materials can combine advantages of different materials.

Hydroxyapatite (HA) is the main inorganic component of human bone tissue[4]. The synthesized HA has good biocompatibility, bone conductivity and strong bonding ability with bone tissue[5]. As an implant material, it can guide the new bone growth and provide a physiological framework for the bone formation[5, 6].

HA and polymer composites have been widely used as bone replacement materials in the world. PLGA, the most widely used of scaffold material in bone and cartilage tissue engineering, has strong mechanical properties and adjustable degradation rate[7]. HA/PLGA composite have the advantages of HA and PLGA, and overcome their disadvantages, such as poor mechanical property, fast degradation rate and weak holding force[8].

At present, adding growth factors, proteins and osteoblasts into the scaffolds mainly improve the biological activity of tissue-engineered scaffolds. However, these additives, inconsistent and unstable in vitro and in vivo, can be easily destroyed by temperature, pH, solvent, enzyme and other factors, which limits their clinical application.

Methyl sulfonyl methane (MSM) is a stable amphiphilic small molecule, which is necessary for the synthesis of collagen[9]. While collagen, necessary for chondrocytes and osteoblasts, can promote the adhesion and migration of bone cells, the formation of collagen matrix, and the reconstruction of bone tissue[10, 11]. However, MSM is easily metabolized and excreted, which lead to short duration of efficacy[12-14].

By introducing bioactive MSM into porous scaffolds, on the one hand, it can improve the biocompatibility of scaffolds; on the other hand, it can prolong the efficacy time by releasing slowly through scaffolds.

In this study, we used the combination of HA and PLGA in the construction of polymer scaffolds. MSM was introduced into polymer scaffolds by freezing phase separation method to prepare five porous scaffolds, named as HA/PLGA porous scaffold, MSM_0.01%_/HA/PLGA porous scaffold, MSM_0.1%_/HA/PLGA porous scaffold, MSM_1%_/HA/PLGA porous scaffold and MSM_10%_/HA/PLGA porous scaffold. The osteoblasts were isolated and cultured in vitro, and the purified osteoblasts were co-cultured with five porous scaffolds to construct the tissue-engineered scaffolds. Meanwhile, a model of rabbit radius defect was constructed to evaluate the biological prospect of five porous scaffolds, which provided experimental basis for the application of the porous scaffolds in bone tissue engineering. The Schematic diagram of bone defect reconstruction based on the fusion of osteoblasts and MSM/HA/PLGA porous scaffolds was shown in Scheme 1.

**Scheme 1.**
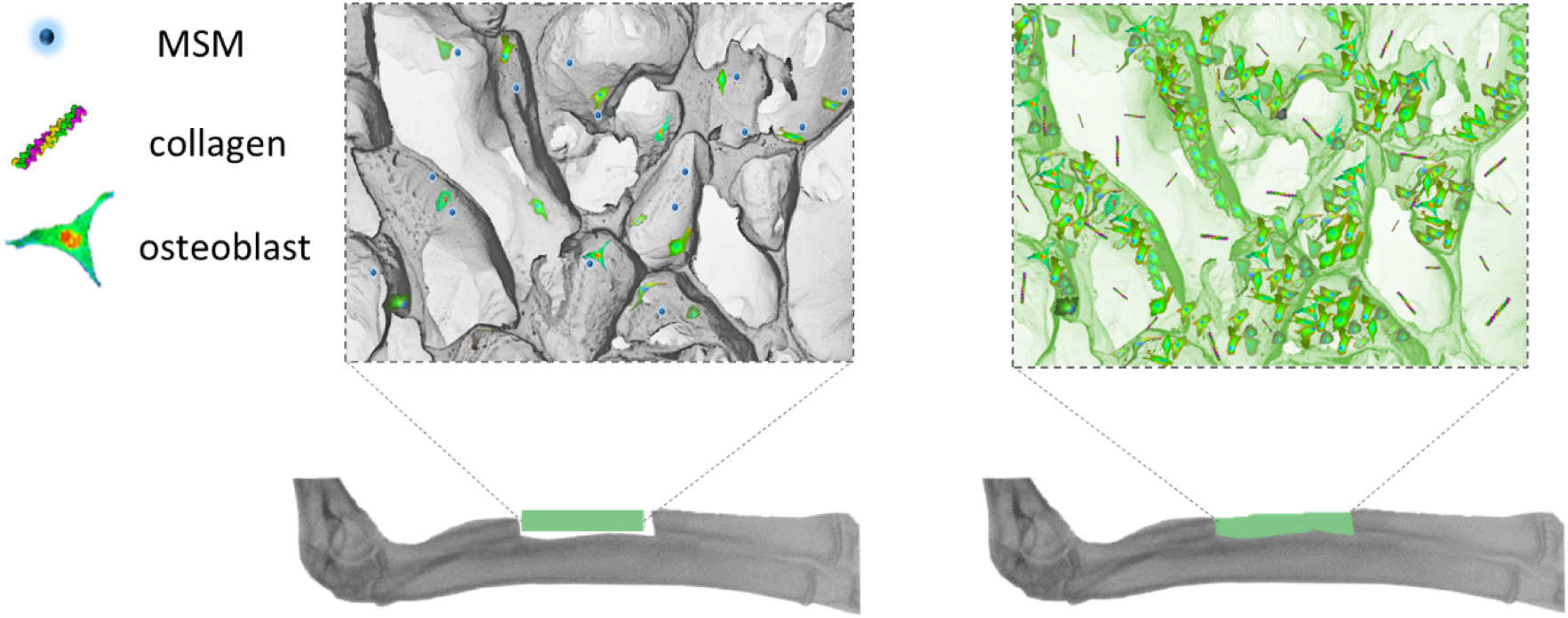
The Schematic diagram of bone defect reconstruction based on the fusion of osteoblasts and MSM/HA/PLGA porous scaffolds.

## Materials and methods

### Experimental materials and reagents

#### Experiments on animals

36 healthy adult white rabbits, male, 2 months old, weighing 2.5 ∼ 3.0 kg, were provided by Experimental Animal Center of JiLin University. All experiments were conducted in accordance with the National Institute of Health Guide for the Care and Use of Laboratory Animals and conducted in line with protocols approved by the Animal Care and Ethics Committee of XuZhou Central Hospital.

#### Reagents and instruments

Main reagents and instruments were listed in the Table 1.

**Table 1.** Main reagents and instruments.

|  |  |
| --- | --- |
| Hydroxyapatite | Changchun Institute of Applied Chemistry<br>Chinese Academy of Sciences |
| Poly (lactic- co - glycolic acid)<br>PLGA (MW. ~10000) | Changchun Institute of Applied Chemistry<br>Chinese Academy of Sciences |
| Dimethyl sulfone | PanJin Far East JinXing Chemical CO.,<br>LTD., China |
| 1,4-dioxane | Sinopharm Chemical Reagent Beijing Co.,<br>Ltd |
| Acetone | Sinopharm Chemical Reagent Beijing Co.,<br>Ltd |
| Dulbecco's modified eagle medium | Gibco |
| Fetal bovine serum | Gibco |
| Trypsin | Gibco |
| MTT | Solarbio |
| DAPI | Sigma |
| Alkaline phosphatase kit | Sigma |
| Penicillin | North China Pharmaceutical Co., Ltd |
| Streptomycin | North China Pharmaceutical Co., Ltd |
| Fluorescence inversion microscope system | Nikon Eclips TE2000-U |
| Digital camera system | Nikon DXM1200F |
| CO<sub>2</sub> incubators | SANYO MCO-18AC |
| Automatic enzyme marker | Multiskan MK3 |
| Vacuum oven | Shanghai Deying Vacuum & Lighting<br>Equipment Co.,Ltd. |
| Material fatigue circulation system | Instron 1121 |
| Field emission scanning electron microscope (FESEM) | Philips XL 30 FEG |
| Inductive Coupled Plasma emission spectrometer (ICP-OES) | Thermo scientific. ( ICAP6300 ) |
| Contact angle meter | Krüss , DSA 10 |

## Methods

### Preparation of MSM/HA/PLGA porous scaffolds

PLGA and MSM were dissolved in 1, 4-dioxane and acetone solutions (10 wt %) at room temperature (25°C). At the same time, HA was dispersed into 1, 4-dioxane solution (10 wt %), treated with ultrasonic treatment for 30 minutes. Mixed three solutions above to prepare the mixture of porous scaffolds, in which the proportion of MSM to MSM/HA/PLGA is MSM: HA: PLGA= (0, 0.01%, 0.1%, 1%, 10%):1:9. MSM/HA/PLGA porous scaffolds were prepared by vacuum freeze-drying for 2 weeks, and were kept in vacuum drying box before used.

### Porosity

The porosity of porous scaffolds was detected by anhydrous ethanol substitution method. Cut the porous scaffolds into strips of 20mm × 5mm × 3mm and weighed them (denoted as W_1_). The porous scaffolds were immersed in a measuring cylinder, containing 4mL of anhydrous ethanol, standing for a few minutes. The measuring cylinder was kept in a vacuum until bubbles were not visible. Recorded the volume at this time as V_1_. Then the porous scaffold was taken out and weighed as W_2_ after drying. The remaining volume of ethanol in the measuring cylinder was recorded as V_2_. The porosity of porous was calculated by formula (1). (ρ_ethanol_= 0.789g/cm^3^)

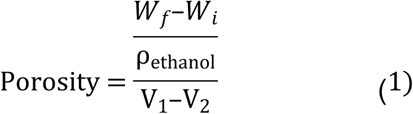

### Contact Angle

The porous scaffolds were dissolved in 1:1 chloroform acetone solution, uniformly suspended, coated on the glass slide, and vacuum-dried for 24 hours. 2 μL of deionized water was directly dropped onto the surface of the sample at room temperature. The Angle θ at which the water drops were tangent to the film on the glass slide were determined and next recorded, three times repeated.

### Mechanical Properties

The compressive and flexural strength of porous scaffolds was tested by Material fatigue circulation system, according to the standards GB/T 1041-1992 and GB/T 16419-1996. The materials were cut into strips of 30mm×5mm×4mm and 10mm×5mm×4mm. The strips were then tested with the loading speed was 5mm/min and 2mm/min respectively. The loading of the strips as they deform was recorded, three times repeated.

### Drug loading efficiency

20mg of each sample was dissolved in 10mL HNO_3_ (1 mol/L), and the volume was fixed to 25mL with deionized water. The content of MSM in solution was measured by ICP-OES, three times repeated.

### Drug release

0.2g of each sample was dispersed into 5mL 0.02M PBS (pH=7.2), and loaded into a dialysis bag. The dialysis bag was then immersed in 10mL PBS solution, with stirring (120 RPM), and a small quantity of PBS solution was taken from the dialysis solution at intervals. The drug release of MSM was calculated by the content of sulfur elements, which was measured by ICP-OES. Three times repeated.

### FESEM characterization

The porous scaffolds were frozen with liquid nitrogen and broken rapidly to obtain brittle sections. The surface morphology of porous scaffolds was observed by FESEM (20KV) after uniform gold spraying.

### Extraction of osteoblasts from rabbits

Primary osteoblasts of rabbits were extracted as follow steps. The newborn milking rabbits were sterilized and killed under aseptic conditions. Cranial parietal was isolated and repeatedly washed with PBS to remove the bloody components. Then the periosteum and soft tissue were removed and the bone fragments were cut into 1mm x 3mm pieces. Bone pieces, digested with 0.25% trypsin for 10 minutes, were transferred to complete culture medium (DMEM, 10% FBS) with evenly distributed, and cultured in an incubator (37°C and 5% CO_2_) for two days. Replaced the culture medium to fresh every two days.

### Purification of osteoblasts

Osteoblasts were purified by differential adhesion as follow steps. Overgrown osteoblasts were cultured by cell passage, and the osteoblasts suspension was stood for 10 min. After then, the supernatant was gently transferred to the new petri dish to keep fibroblasts from mixing. Repeated the procedure three times to remove the fibroblasts thoroughly.

### Osteoblast implantation

The porous scaffolds were cut into cylindrical shapes (11mm in diameter and 5mm in height), sterilization by dry heat for 60 minutes. The cylindrical samples were moved in 24-well plates, and were prewetted with culture medium in the incubator for 24 hours. The third generation of rabbit osteoblasts were digested with 0.25% trypsin and were seeded in the scaffold with a density of 10.0×10^4^ cells /well, incubated at 37°C and 5% CO_2_ for 4 hours, so that the osteoblasts could better adhere to the porous scaffolds well. Changed the culture medium every two days.

### Cell adhesion permeability

After continuous cultivation of osteoblasts on porous scaffolds for 7 days, the medium was removed. The scaffolds were then fixed with glutaraldehyde (2.5%) for 15 minutes and washed with PBS three times later. After that, the porous scaffolds were dyed with DAPI for 10 minutes and washed with PBS three times. The adhesion and permeability of rabbit osteoblasts on porous scaffolds were evaluated by fluorescence microscopy.

### Proliferation of osteoblasts

Osteoblasts were cultured on porous scaffolds for 3d, 7d and 14d, and 100μL/well MTT solution was added to 24-well plates for 4h. The medium was sucked out and 750μL (0.4 mol/L) isopropanol hydrochloride solution was added to dissolve the porous scaffold for 15min. After the mixture was evenly dispersed, 200μL mixture was transferred from 24-well plate to the 96-well plate. The light absorption value of each well was detected by automatic microplate reader at 540nm, three times repeated.

### Activity of alkaline phosphatase

After continuous cultivation of osteoblasts on the porous scaffolds for 7 days, the culture medium was removed and the leftover was washed with PBS three times. Trypsin was used for digestion for 3-5min, and the complete culture medium (DMEM, 10% FBS) was used to terminate digestion. After then, the supernatant was removed by centrifugation (1500 RPM, 5min), and the cells were collected and lysed with 0.2% NP-40 cell lysate LML. Ultrasonic cracking was performed in ice bath for 3-5min, and the suspension was centrifuged again (1500 RPM, 5min). The supernatant was taken and the activity of ALP was determined according to the instructions of the ALP activity detection kit.

### Establishment of animal model of radius defect

The weight of the white rabbits was weighed, and the rabbits were anesthetized with Sumianxin? at the dose of 0.2ml/kg. After anesthesia, the hair of the forelimbs was shaved and the body surface of the forelimbs was cleaned. The rabbits were fixed on the laboratory bench, with their limbs fully abducted. The limbs were sterilized with iodine and the rabbits were placed on the sterile sheets. An incision of about 2.5cm long was made lengthwise along the middle and lower radius of the forearm. Subcutaneous tissue, fascia and muscle were then separated layer by layer to expose the radius. A segmental bone defect of 20mm was made in the middle and lower radius with an electric saw.

### Tissue-engineering scaffolds implantation

Porous scaffolds were prepared according to the radius diameter of rabbits, and seeded with osteoblasts for two days to conduct the tissue-engineering scaffolds. The tissue-engineering scaffolds were implanted into the radial defects separately, while no disposal was required in the blank control group. Then, the surgical wound was rinsed with gentamicin and sutured layer by layer. Finally, the skin was Sutured and sterilized with iodine. Intramuscular injections of 400,000 units of penicillin were given daily for one week. X-ray examination was performed to evaluate the treatment effect of scaffold implantation for radius defect.

## Results and discussion

The surface morphology of the scaffolds was obtained by FESEM, as shown in Figure 1(A). MSM _(0%, 0.01%, 0.1%, 1%, and 10%)_/HA/PLGA porous scaffolds had a typical honeycomb pore structure. When no MSM was mixed into the HA/PLGA scaffold, the pores were evenly distributed and the pore walls were clear and smooth. When the MSM content was low, the MSM _(0.01%, 0.1%)_/HA/PLGA scaffold had a similar pore structure with HA/PLGA scaffold. A few of small holes appeared on the wall of the big holes, which were interconnected through small holes. However, as the content of MSM further increased, the pore structure gradually loosen. When the content of MSM rose to 1%, the pore of scaffolds fractured with irregular structure.

**Figure 1.**
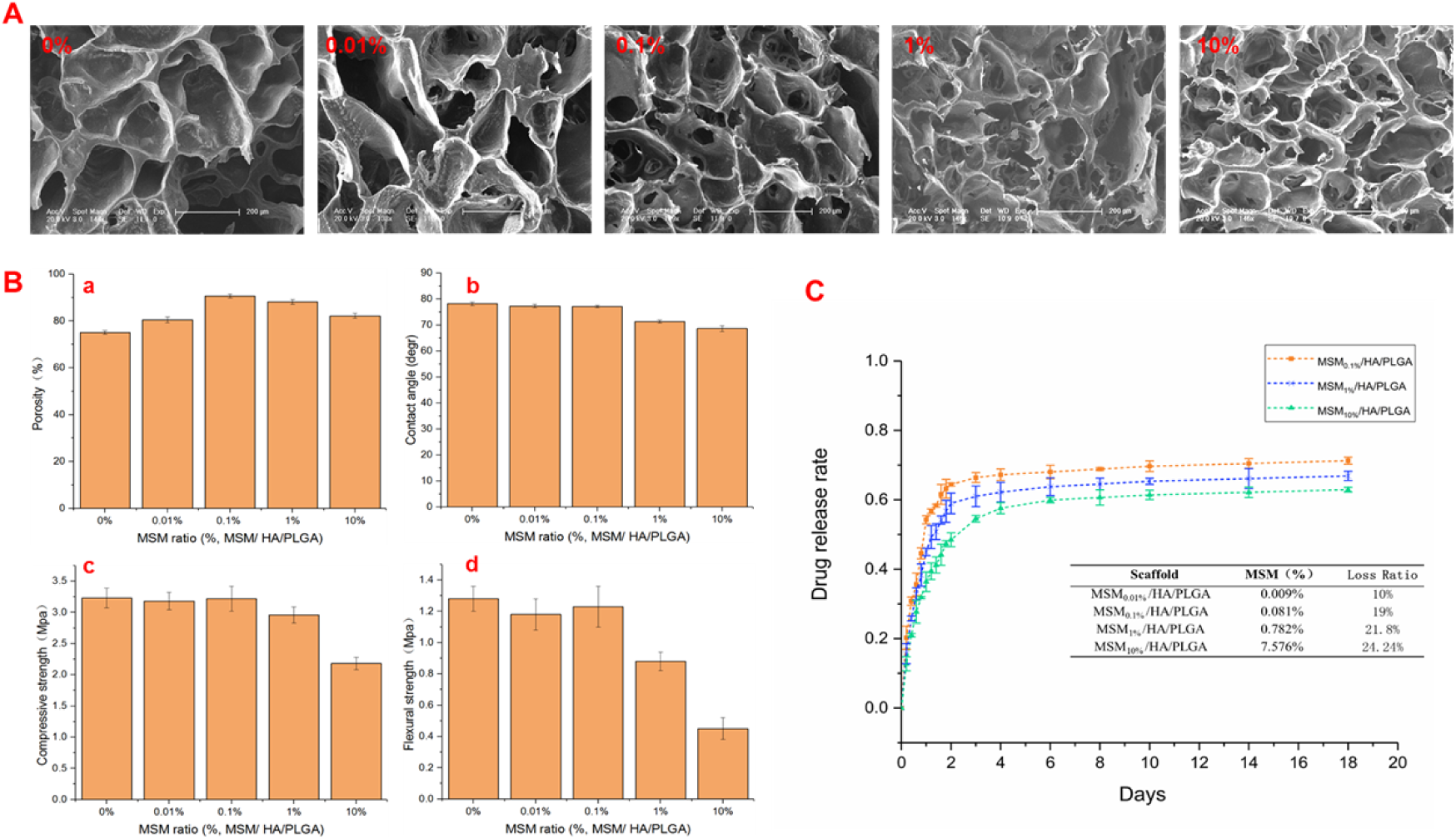
(A) SEM characterization of MSM/HA/PLGA scaffolds with different proportion of MSM, Scaler: 200 μm; (B) The physical properties(1, Contact angle; 2, Flexural strength; 3, Porosity; 4, Compressive strength) of scaffolds with different proportion of MSM; (C) The line chart of MSM release rate of scaffolds with different proportion of MSM, Insert table: The actual content and the loss ratio of MSM in MSM _(0.01%, 0.1%, 1% and 10%)_/HA/PLGA porous scaffolds

When the content of MSM reached 10%, the pore wall became thinner and small pores gradually expanded until the difference between the larger pores was no longer obvious. With the increase of MSM content, the surface of porous scaffolds became more and more rough, which was conducive to the crawling adhesion of cells.

As can be seen from Figure 1 (B-a), there is no significant difference in the porosity of different porous scaffolds. When the MSM content is low (0%, 0.01%, and 0.1%), the porosity gradually increased with the increase of MSM content, and when the MSM content was 0.1%, the porosity reached the highest 90.6%. However, with the further increase of MSM content, the porosity of the scaffold decreased.

The contact Angle θ of scaffolds fully demonstrates the hydrophilicity of the material. Generally speaking, if θ <90 °, the solid surface is hydrophilic. The smaller the Angle, the better the wettability. If θ > 90 °, the solid surface is hydrophobic, that is, the liquid does not easily wet the solid and moves easily over the surface.

As shown in Figure 1(B-b), the contact Angle of MSM_(0%, 0.01%, 0.1%, 1% and 10%)_/HA/PLGA porous scaffolds decreased with the increase of MSM content, indicating that the porous scaffolds are more hydrophilic with the increase of MSM content, possibly because MSM is a water-soluble compound.

The mechanical properties were analyzed by testing the compressive strength and flexural strength of scaffold materials, as shown in Figure 1 (B-c) and Figure 1 (B-d). HA/PLGA scaffold had the highest Compression Strength (3.23MPa) and Flexural Strength (1.28MPa) and the mechanical properties of the MSM/HA/PLGA scaffolds did not decrease significantly when the content of MSM was less than 0.1%. However, when the MSM content was higher than 0.1%, the mechanical properties of scaffolds decreased significantly, which due to the excessive MSM content, leading to the decline of polymer forming ability and the destruction of pore structure.

The actual content of MSM in porous scaffolds was determined by ICP-OES, as shown in the insert table of Figure 1 (C). The results showed that the actual content of MSM in porous scaffolds was little different from the theoretical content. For MSM _(0.01%)_/HA/PLGA porous scaffolds with relatively complete pore structure, the loss rate of MSM in the preparation of scaffolds was only 10%. At the same time, the loss ratio of MSM in MSM _(0.1%, 1%, and 10%)_/HA/PLGA porous scaffolds reached around 20% of the theoretical value, which was caused by the great hydrophilicity of MSM and the incomplete pore structure.

The release rate of MSM of porous scaffolds was calculated from the theoretical value and the actual value, which measured by ICP-OES in 18 days, as shown in Figure 1 (C).

The results showed that the release rate of all porous scaffolds could finally reach 60% of the theoretical value, while the lower the MSM content of porous scaffold had, the higher the release rate was. For all porous scaffold, MSM suddenly released in the first two days, and slowly released in the following days. This is because MSM is a water-soluble molecular compound, and MSM/ HA/PLGA porous scaffolds have a large number of pores, which allow solution to fully contact with MSM. Two days later, the release of MSM entered into a slow release period, which indicated that porous scaffolds could prolong the release time of MSM and thus the efficacy time of MSM. As can be seen from Figure 2, the cell adhesion permeability showed the number of cells on MSM _(0.01%,0.1% and 1%)_ /HA/PLGA porous scaffolds was significantly higher than that of HA/PLGA and MSM_10%_ /HA/PLGA scaffolds, which was also confirmed by cell proliferation efficiency and ALP activity.

**Figure 2.**
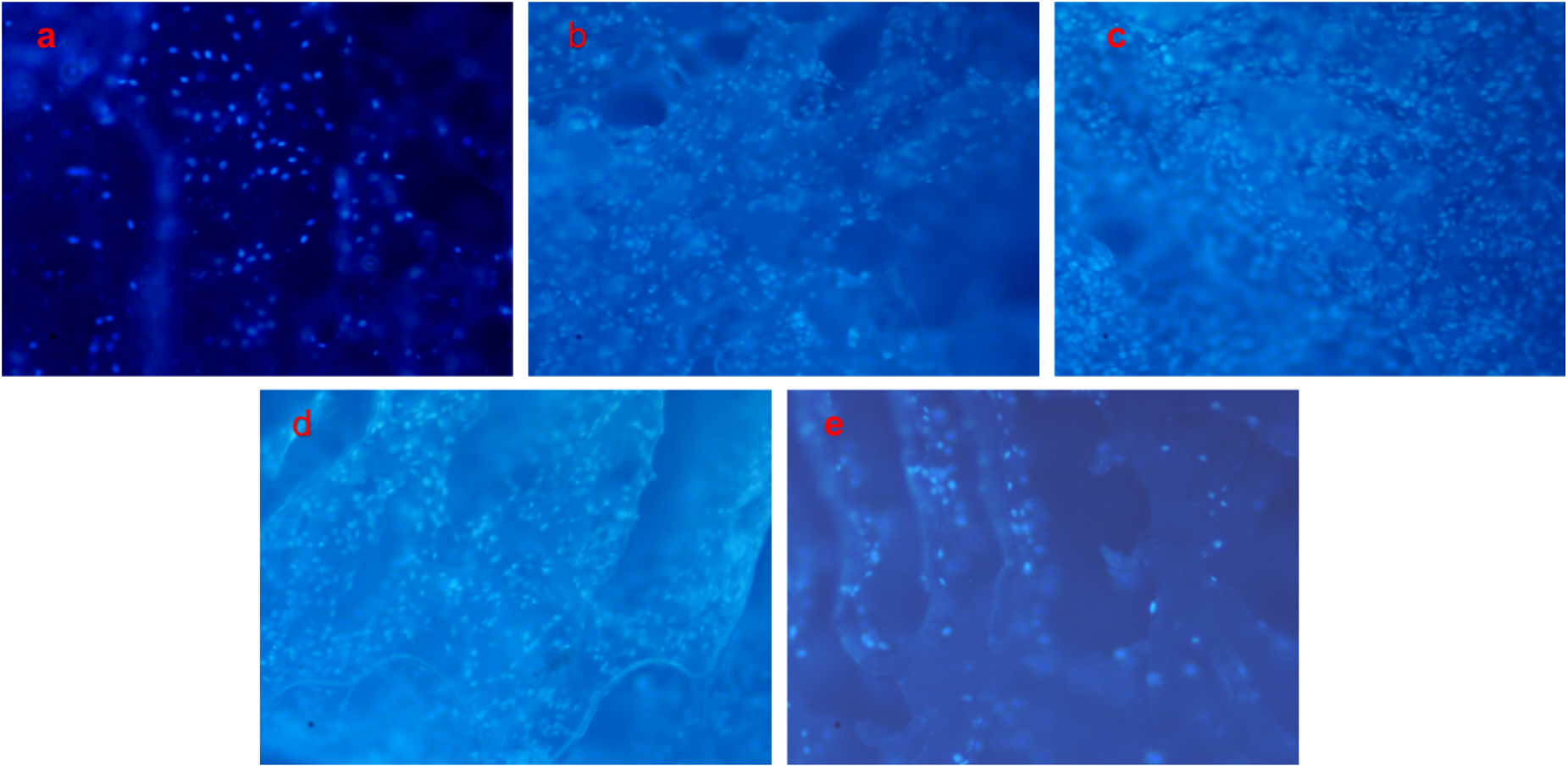
Fluorescence image of osteoblasts on different scaffolds after DAPI staining: HA/PLGA(a), MSM_0.01%_/HA/PLGA(b), MSM_0.1%_/HA/PLGA(c), MSM_1%_/HA/PLGA(d) and MSM_10%_/HA/PLGA(e)

At the same time, the results showed that osteoblasts adhered tightly to the surface of the porous scaffolds, and were able to proliferate, migrate and infiltrate into the scaffold material, so as to evenly distribute throughout the scaffolds.

The uniform distribution of osteoblasts in the whole scaffold benefits from the good biocompatibility of porous scaffolds, the pore size suitable for osteoblasts’ migration and the good connectivity between pores.

As can be seen in Figure 3, after continuous cultivation of osteoblasts on porous scaffolds for 3days, 7 days and 14 days, the proliferation activity and cell number of osteoblasts increased obviously. Osteoblasts proliferation on MSM _(0.01%, 0.1%, 1% and 10%)_/HA/PLGA porous scaffolds was significantly better than that on HA/PLGA scaffold in all periods. Besides, the content of MSM in scaffolds had a great influence on the proliferation of osteoblasts. When the content of MSM in porous scaffolds was less than 0.1%, the number of osteoblast on the scaffolds increased with the increase of MSM content. However, when the MSM content in porous scaffolds was higher than 0.1%, the number of osteoblasts on the scaffolds slightly decreased. There was little difference between the number of osteoblasts on the MSM_1%_/HA/PLGA and MSM_10%_/HA/PLGA scaffolds.

**Figure 3.**
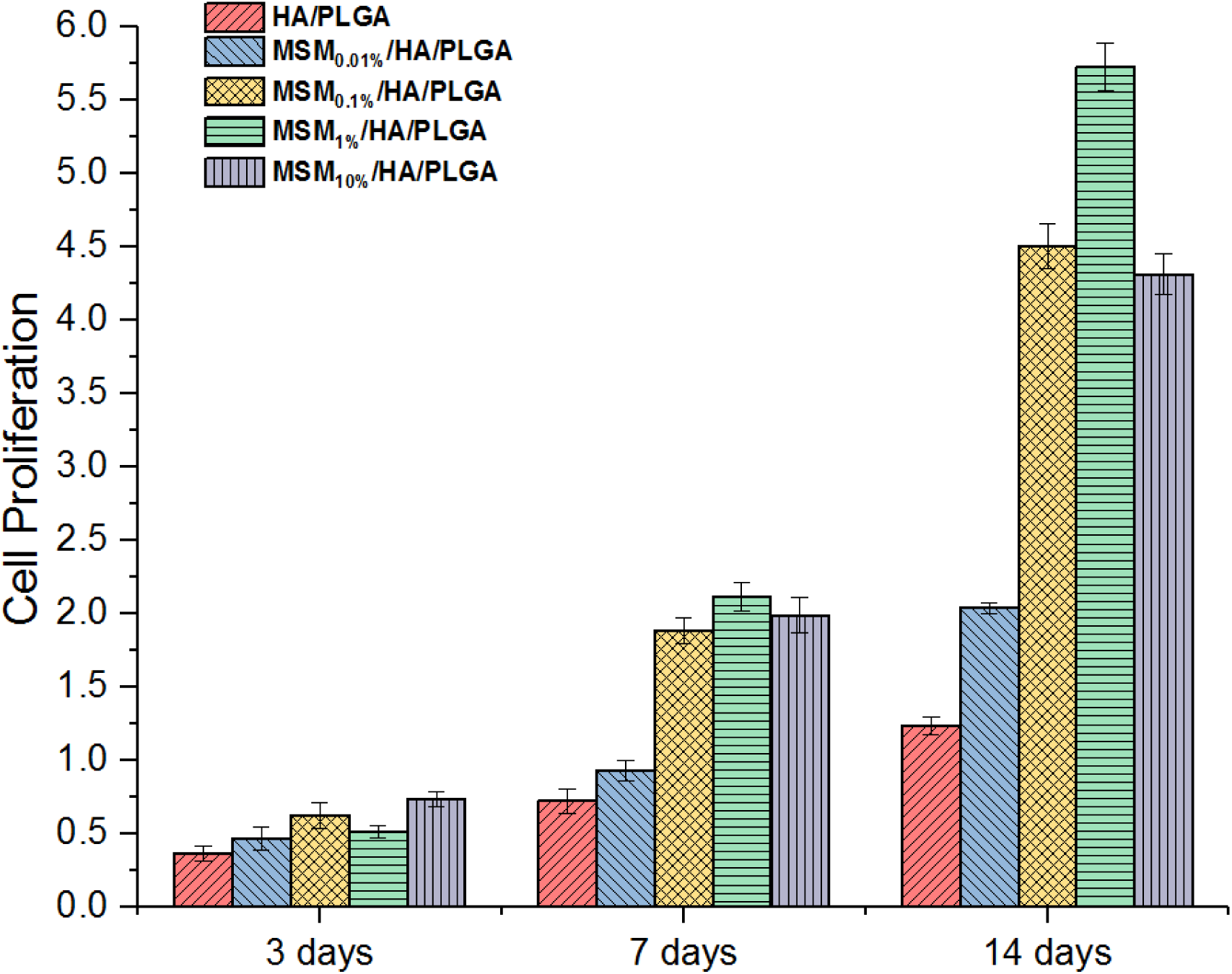
Proliferation of osteoblasts on different scaffolds for 3 days, 7 days and 14 days.

We speculated that there are several reasons for this proliferation. 1) MSM, a bioactive small molecule, is essential for the synthesis of collagen, which is a major component of chondrocytes and bone extracellular matrix. 2) The addition of MSM increases the surface roughness of porous scaffold, which is conducive to cell adhesion. 3) Excessive addition of MSM destroys the pore structure of porous scaffolds, thus reducing the area where cells can adhere. 4) Excessive addition of MSM also leads to excessive concentration of MSM in the scaffold, which hinders the normal proliferation of osteoblasts.

Similar effects were found in the difference in alkaline phosphatase activity of osteoblasts on porous scaffolds for 7 days, as shown in Figure 4.

Alkaline phosphatase (ALP) is an extracellular enzyme of osteoblasts, and its expression activity is an obvious feature of osteoblasts differentiation. The main physiological function of ALP in the body is to hydrolyze phosphate ester in the process of osteogenesis to provide necessary phosphoric acid for the deposition of hydroxyapatite. At the same time, ALP hydrolyzes pyrophosphate to remove the inhibition of pyrophosphate on bone formation, which is conducive to osteogenesis. In the most study of bone tissue engineering, ALP activity is used as an important index to reflect the biological activity of bone repairing materials.

**Figure 4.**
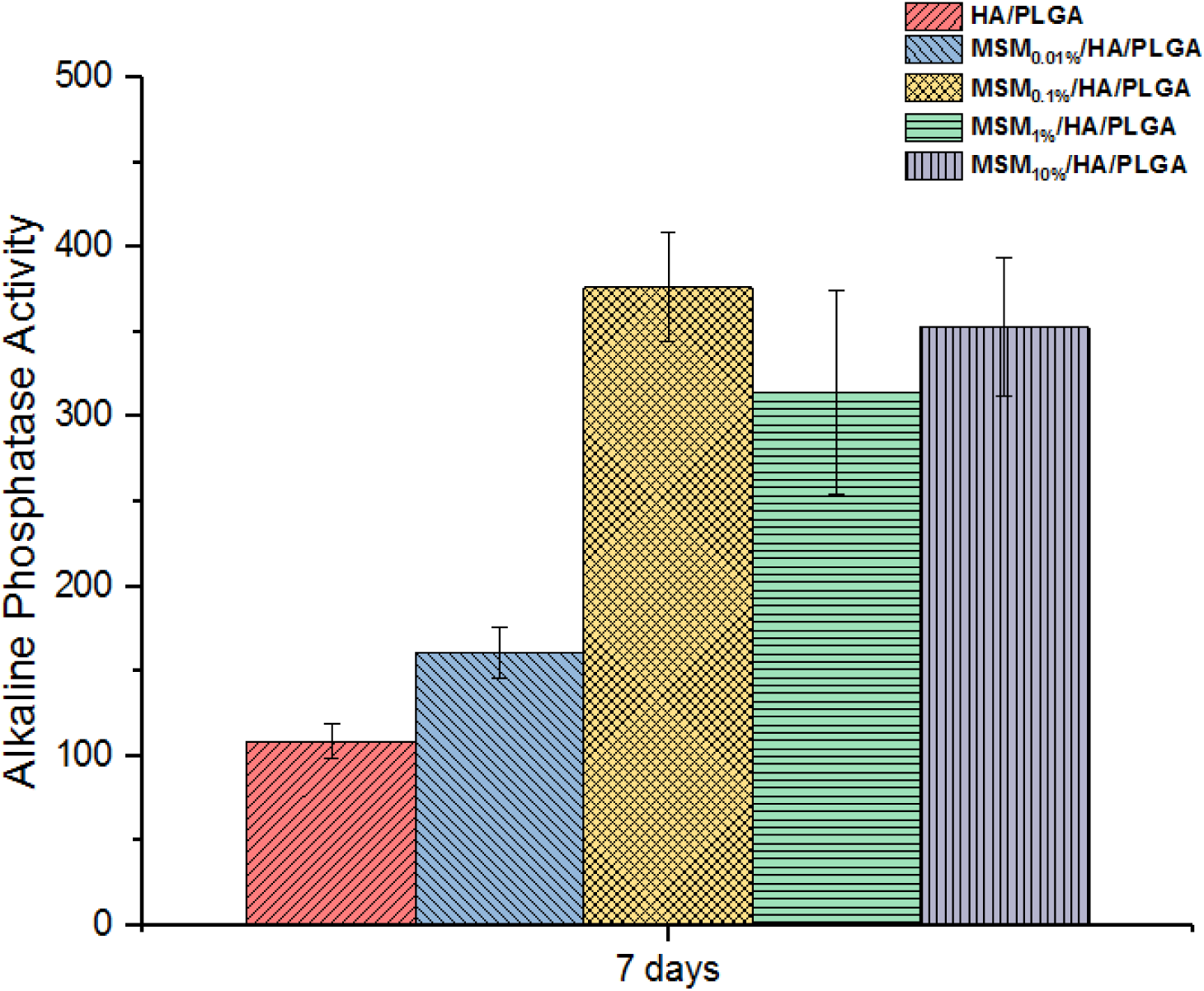
Alkaline phosphatase activity of osteoblasts on different scaffolds.

When MSM content in the composite scaffolds was less than 0.1%, ALP activity increased with the increase of MSM content. However, when MSM content continued to increase, ALP activity decreased.

We hypothesized that when the MSM content in the composite scaffold was low (0.01%, 0.1%), MSM promoted the synthesis of collagen, which is necessary for the proliferation of osteoblasts, and the proliferation of osteoblasts lead to a significant increase in ALP activity. However, when the content of MSM in the porous scaffold is high (1%, 10%), a large quantity of MSM is released, resulting in high local MSM content and side effects, thus inhibiting the activity of ALP.

Digital Radiographs and morphological observations of rabbit radius defects implanted with the different tissue-engineering scaffolds for 4 weeks and 12 weeks were shown in Figure 5.

4 weeks after surgery: Figure 5(A_1_) showed a few callus growing from the broken bone in the blank control group, while other groups showed significant callus formation and bridged changes. When the tissue-engineering scaffold implanted at the defect bone did not contain MSM, the amount of callus formed was small, and the callus could be seen to crawl along the ulnar side, as shown in Figure 5 (B_1_). When the implanted tissue-engineering scaffold contained MSM _(0.01%, 0.1%, 1% and 10%)_, obvious callus formed at the fracture site, the fracture line was blurred, and the bone connection was initially formed, as shown in Figure 5 (C_1_∼G_1_).

12 weeks after surgery: The bone defect was still obvious in the blank control group, while other groups formed continuous callus connections. Low density shadow can be seen at the far folding end of HA/PLGA scaffold group, while bone marrow cavity and cortical contour, basic reconstruction of bone defects, the disappearance of fracture line can be seen in other tissue-engineering scaffold groups, which containing MEM_(0.01%, 0.1%, 1% and 10%)_.

**Figure 5.**
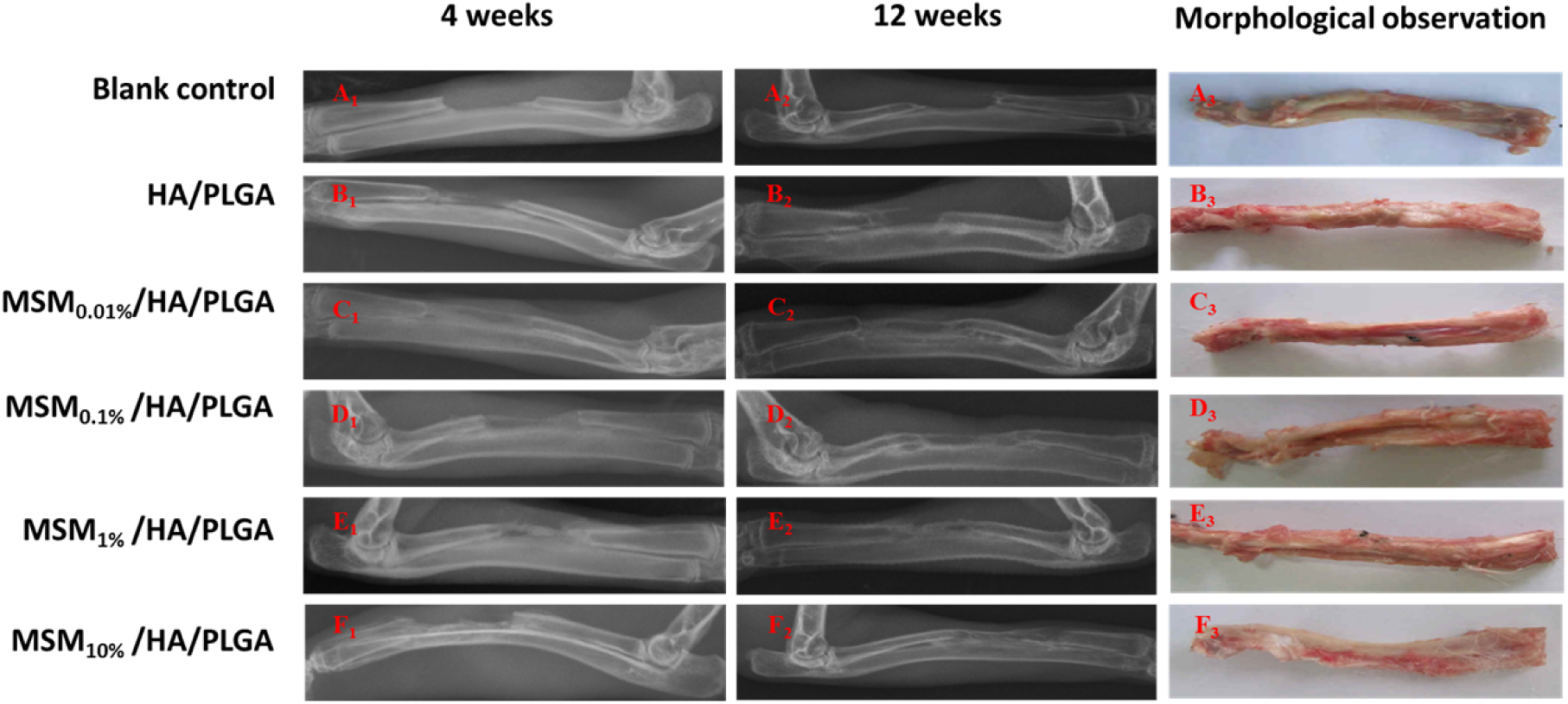
Digital Radiograph(DR) and Morphological observation of rabbit radius defects implanted with the different scaffolds: blank control (A_1-3_), HA/PLGA(B_1-3_), MSM_0.01%_/HA/PLGA(C_1-3_),MSM_0.1%_/HA/PLGA (D_1-3_), MSM_1%_/HA/PLGA (E_1-3_) and MSM_10%_/HA/PLGA (F_1-3_).

The results showed that the MSM _(0%, 0.01%, 0.1%, 1% and 10%)_/ HA/PLGA groups have significantly superior bone reconstruct ability than HA/PLGA group.

This may be because when degraded slowly in vivo, the porous scaffolds continuously released bioactive small molecule MSM, calcium ions and phosphate particles, which can promote the proliferation of bone cells and the secretion of bone matrix. In addition, the porous scaffolds mixed with a certain quantity of MSM can provide the biological activity of polymers, promoted the adhesion and proliferation of osteoblasts, to accelerate bone repair and improve repair efficiency.

Morphological observations of rabbit radius defects implanted with the tissue-engineering scaffolds for 12 weeks showed that the tissue-engineering scaffolds had obvious repair effect on bone defects, while no automatic repair effect observed in the blank control. The tissue-engineering scaffolds, implanted at the site of the defect bone, bridged to the surrounding bone tissue, with no bone malformation, stent displacement or stent fracture being found. No obvious inflammatory reaction and abnormal effusion was found at the site of the tissue-engineering scaffolds implantation.

## Conclusion

In this study, we prepared MSM/HA/PLGA porous scaffolds by freeze-drying method, which has appropriate pore size, high porosity, excellent mechanical properties, and high drug loading rate and control release.

The experimental results of co-incubation of porous scaffolds with osteoblasts showed that osteoblasts could proliferate and migrate on porous scaffolds to cover the whole scaffold. The animal experimental results of tissue-engineering scaffolds, prepared by incubation of MSM _(0%, 0.01%, 0.1%, 1% and 10%)_/ HA/PLGA porous scaffolds and osteoblasts for two days, implanted in the rabbit radius defects showed good bone repair ability.

The difference of porous scaffolds with different MSM content in bone defect reconstruction indicates the great role of MSM in bone repair, which provided experimental basis for the application of the MSM/HA/PLGA porous scaffolds in bone tissue engineering.

## Acknowledgements

This work was supported by the Xuzhou Science and Technology Bureau Project (KC18039), Postdoctoral Funding Project of Jiangsu Province (2018K244C), Double Innovation in Jiangsu Province (2016SC04), Xuzhou Clinical Technical Backbone Research Program (2018GG027).

